# Development of structure-function coupling in human brain networks during youth

**DOI:** 10.1101/729004

**Authors:** Graham L. Baum, Zaixu Cui, David R. Roalf, Rastko Ciric, Richard F. Betzel, Bart Larsen, Matthew Cieslak, Philip A. Cook, Cedric H. Xia, Tyler M. Moore, Kosha Ruparel, Desmond Oathes, Aaron Alexander-Bloch, Russell T. Shinohara, Armin Raznahan, Raquel E. Gur, Ruben C. Gur, Danielle S. Bassett, Theodore D. Satterthwaite

## Abstract

The protracted development of structural and functional brain connectivity within distributed association networks coincides with improvements in higher-order cognitive processes such as working memory. However, it remains unclear how white matter architecture develops during youth to directly support coordinated neural activity. Here, we characterize the development of structure-function coupling using diffusion-weighted imaging and *n*-back fMRI data in a sample of 727 individuals (ages 8-23 years). We found that spatial variability in structure-function coupling aligned with cortical hierarchies of functional specialization and evolutionary expansion. Furthermore, hierarchy-dependent age effects on structure-function coupling localized to transmodal cortex in both cross-sectional data and a subset of participants with longitudinal data (*n*=294). Moreover, structure-function coupling in rostrolateral prefrontal cortex was associated with executive performance, and partially mediated age-related improvements in executive function. Together, these findings delineate a critical dimension of adolescent brain development, whereby the coupling between structural and functional connectivity remodels to support functional specialization and cognition.

## INTRODUCTION

The human cerebral cortex is organized along a functional hierarchy extending from unimodal sensory cortex to transmodal association cortex (1, 2). This macroscale functional hierarchy is anchored by an anatomical backbone of white matter pathways that coordinate synchronized neural activity and cognition. Both primate cortical evolution and human brain development have been characterized by the targeted expansion and remodeling of transmodal association areas (3, 4), which underpin the integration of sensory representations and abstract rules for executing goals. The protracted development of transmodal association cortex in humans provides an extended window for activity-dependent myelination (5) and synaptic pruning (6). This period of cortical plasticity sculpts functional specialization in transmodal association cortex, and may be critical for developing higher-order executive functions such as working memory, mental flexibility, and inhibitory control (7).

Characterizing the functional specialization of cortical areas based on their patterns of connectivity has been central to understanding hierarchies of brain organization (8, 9). Network theory has provided a parsimonious framework for modeling structure-function mappings in neurobiological systems across species and spatial scales (10). Convergent evidence has highlighted the strong correspondence between measures of structural and functional brain connectivity at different spatiotemporal scales, from neural populations (11) to specialized cortical regions (12), and large-scale brain networks (13–15). However, only sparse data exists regarding how the maturation of white matter architecture during human brain development supports coordinated fluctuations in neural activity underlying cognition. Furthermore, aberrant development of structural constraints on functional communication could contribute to deficits in executive function and the emergence of neuropsychiatric disorders during adolescence (16, 17).

Structure-function coupling describes structural support for functional communication, and occurs when a cortical region’s profile of inter-regional white matter connectivity predicts the strength of inter-regional functional connectivity. Here, we describe the cortical topography of structure-function coupling and delineate how it evolves with development. To do this, we tested three related hypotheses. First, we hypothesized that structure-function coupling would reflect the functional specialization of a cortical area. Specifically, we predicted structure-function coupling would be high in somato-sensory cortex, due to highly conserved programming that governs the early development of specialized sensory hierarchies (18). Conversely, we expected that structure-function coupling would be low in transmodal association cortex, where functional communication may have become untethered from genetic and anatomical constraints through rapid evolutionary expansion (18). Second, based on evidence of prolonged activity-dependent myelination during development (5), we hypothesized that developmental increases in structure function-coupling would be localized to transmodal association cortex. Third and finally, we hypothesized that this structure-function coupling in transmodal cortex would predict individual differences in executive functioning.

## RESULTS

To characterize the development of structure-function coupling in youth, we quantified the degree to which a brain region’s structural connections support coordinated fluctuations in neural activity. Leveraging multi-modal neuroimaging data from 727 participants ages 8-23 years old, we applied probabilistic diffusion tractography and estimated functional connectivity between each pair of cortical regions during a fractal *n*-back working memory task. While intrinsic functional connectivity estimated at rest reflects spontaneous fluctuations in neural activity during unconstrained cognitive states, functional connectivity measured during a working memory task can amplify individual differences in neural circuitry underlying executive performance (19). For each participant, two 400 × 400 weighted adjacency matrices encoding the structural and functional connectome, respectively, were constructed using the same cortical parcellation (20). Structure-function coupling was measured as the Spearman rank correlation between regional structural and functional connectivity profiles (**Fig. 1**).

**Figure 1.**
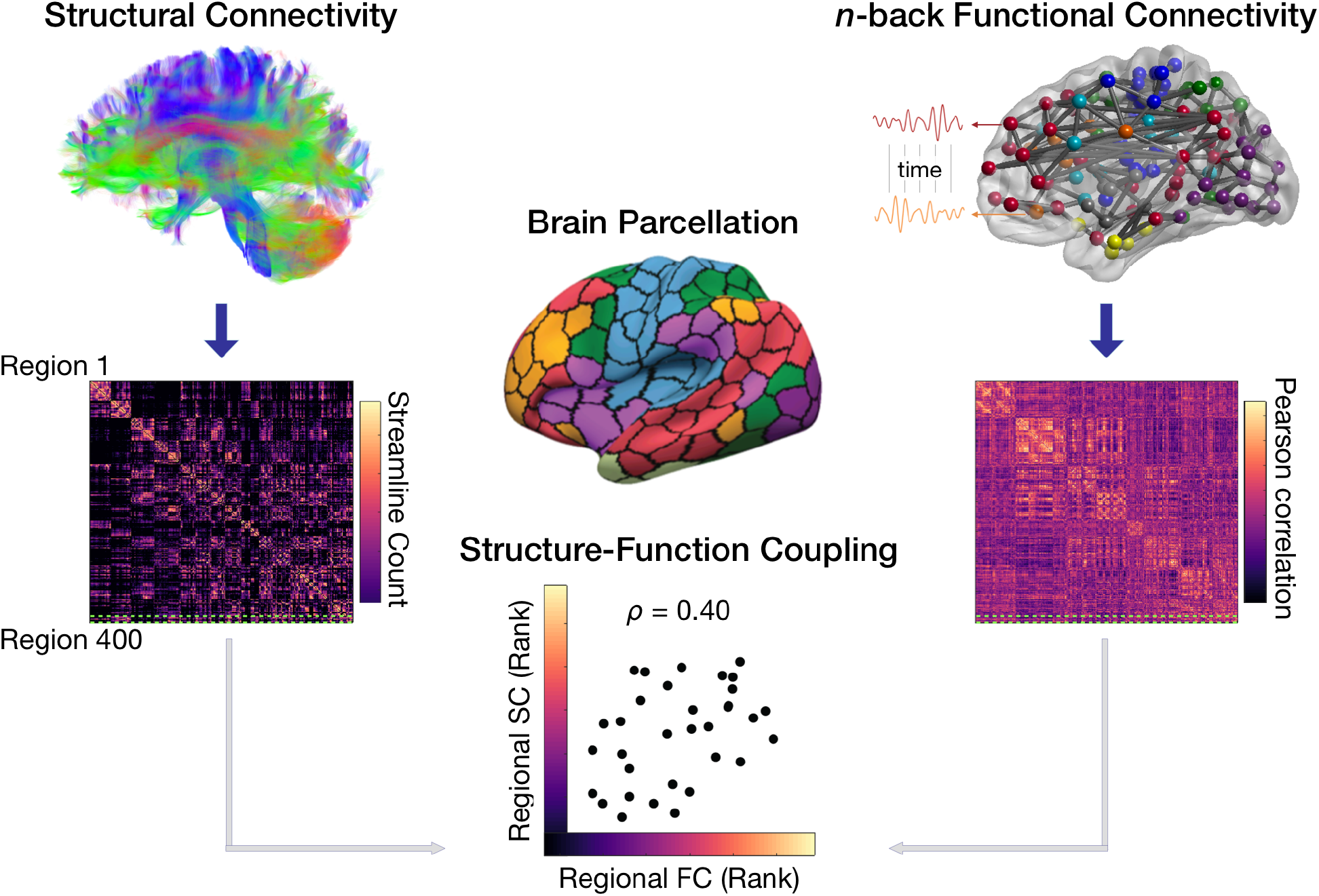
Measuring structure-function coupling in human brain networks. Nodes in structural and functional brain networks were defined using a 400-region cortical parcellation based on functional homogeneity in fMRI data (20). For each participant, regional connectivity profiles were extracted from each row of the structural or functional connectivity matrix, and represented as vectors of connectivity strength from a single network node to all other nodes in the network. Structure-function coupling was then measured as the Spearman rank correlation between nonzero elements of regional structural and functional connectivity profiles.

### Variability in structure-function coupling reflects gradients of functional specialization

As a first step, we assessed whether the spatial distribution of structure-function coupling aligns with fundamental properties of cortical organization. The spatial correspondence between structure-function coupling and other cortical properties was assessed using a conservative spatial permutation test, which generates a null distribution of randomly rotated brain maps that preserve the spatial covariance structure of the original data (21) (denoted *p_spin_*). Notably, the coupling between regional structural and functional connectivity profiles varied widely across the cortex (**Fig. 2A**), with higher coupling in primary sensory and medial prefrontal cortex compared to lateral temporal and frontoparietal regions with lower coupling. To assess the relationship between structure-function coupling and functional specialization, we calculated the participation coefficient, a graph measure that quantifies the diversity of connectivity across functionally specialized modules (22). Brain network nodes with a high participation coefficient exhibit diverse inter-modular connectivity, thereby having the capacity to integrate information across distinct brain modules, while nodes with a low participation coefficient exhibit more locally segregated connectivity within that node’s module. Variability in structure-function coupling was significantly associated with the participation coefficient, calculated for both structural (*r*=−0.28, *p_spin_*=0.001; **Fig. 2B**) and functional (*r*=−0.17, *p_spin_*=0.037; **Fig. 2C**) brain networks. Brain regions exhibiting relatively high structure-function coupling were localized in segregated regions of primary sensory and medial prefrontal cortex, while regions with diverse inter-modular connectivity had relatively lower structure-function coupling.

**Figure 2.**
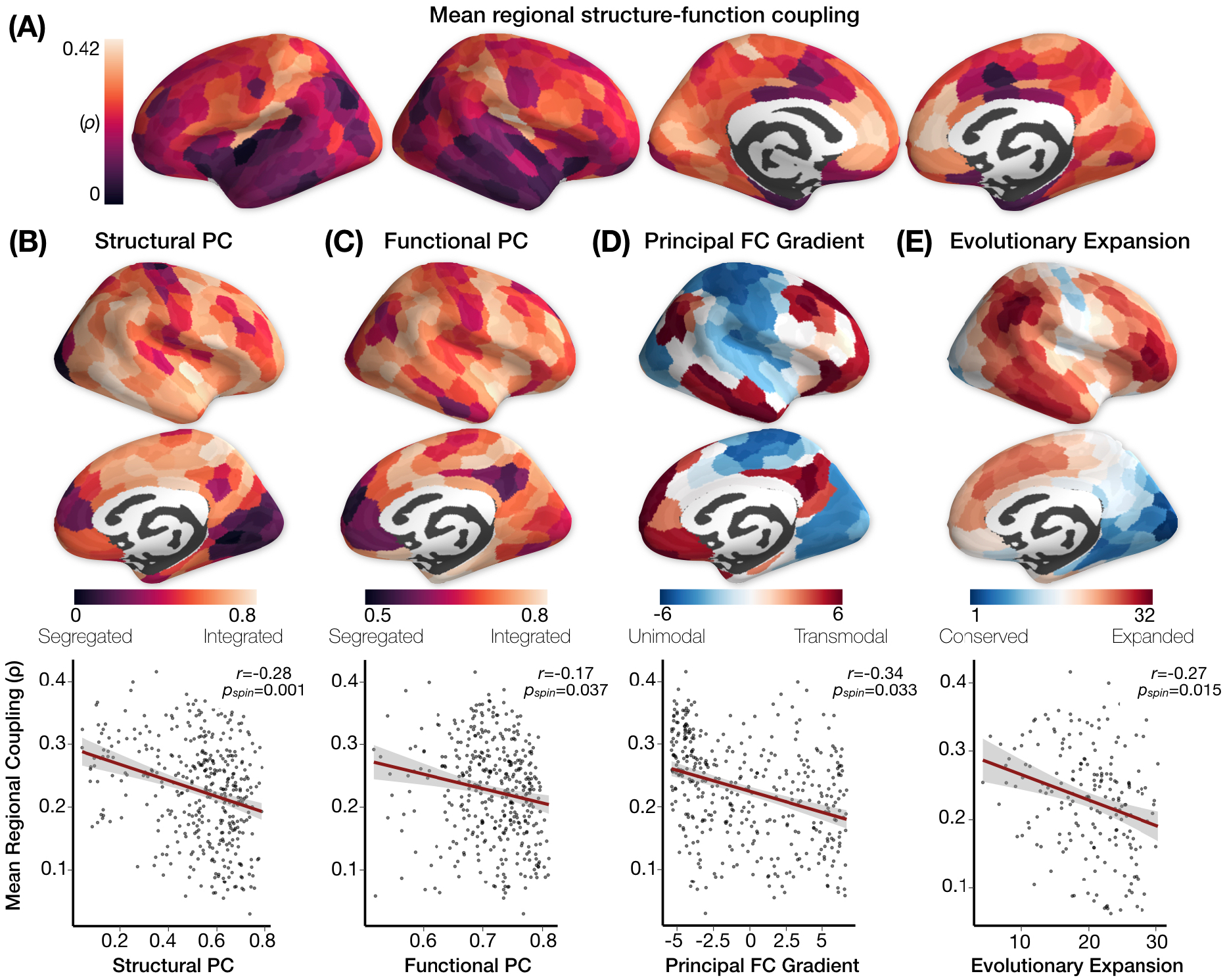
Variability in structure-function coupling reflects cortical hierarchies of functional specialization. The coupling between regional structural and functional connectivity profiles during the *n*-back working memory task varied widely across the cortex. (**A**) Primary sensory and medial prefrontal cortex exhibited relatively high structure-function coupling, while lateral temporal and frontoparietal regions had relatively low coupling. (**B**) Structure-function coupling was significantly associated with the structural participation coefficient (PC), and (**C**) the functional participation coefficient, a measure of the diversity of inter-module connectivity. (**D**) Variability in structure-function coupling also reflected a brain region’s position along the macroscale functional gradient from unimodal to transmodal processing, and (**E**) recapitulated patterns of evolutionary expansion in cortical surface area from macaques to humans. The significance of regional correlations was evaluated using non-parametric spatial permutation testing (denoted *p_spin_*).

Next, we evaluated whether variability in structure-function coupling reflects a macroscale functional hierarchy defined using an independent data-set (2), which captures a primary dimension of variance in intrinsic functional connectivity from unimodal sensory areas to transmodal association cortex. Structure-function coupling aligned significantly with the principal gradient of functional connectivity: unimodal sensory regions exhibited relatively strong structure-function coupling, while transmodal regions at the apex of the functional hierarchy exhibited weaker coupling (*r*=−0.34, *p_spin_*=0.033; **Fig. 2D**). We also tested the hypothesis that functionally specialized somatosensory cortex with evolutionarily conserved organization would exhibit strong structure-function coupling, while highly expanded transmodal cortex would exhibit relatively low structure-function coupling to facilitate functional diversity and cognitive flexibility. Our results were consistent with such an account, as structure-function coupling was significantly correlated with evolutionary expansion of cortical surface area (*r*=−0.27, *p_spin_*=0.015; **Fig. 2E**). Highly conserved sensory areas had relatively strong structure-function coupling, while highly expanded transmodal areas had relatively weak coupling. Together, our results demonstrate that structure-function coupling reflects cortical hierarchies of functional specialization and evolutionary expansion.

### Hierarchy-dependent development of structure-function coupling

While previous work has largely focused on global relationships between group-averaged structural and functional brain networks in adults, here we sought to understand how regional structure-function coupling develops from childhood through adulthood. Regional associations between structure-function coupling and age were assessed using generalized additive models (GAM) with penalized splines, including sex and in-scanner head motion as additional covariates. Age-related differences in structure-function coupling were broadly distributed across lateral temporal, inferior parietal, and prefrontal cortex (**Fig. 3A**). Notably, age-related increases in coupling were disproportionately enriched within a unique subset of functionally segregated areas of the default mode network (*F*=12.54, *p*<10^−10^); **Fig. 3B**). Moreover, the magnitude of age-related differences in structure-function coupling was significantly correlated with the functional participation coefficient (*r*=−0.19, *p_spin_*=0.013; **Fig. 3C**), and the functional gradient from unimodal to transmodal processing (*r*=0.28, *p_spin_*=0.009; **Fig. 3D**). The spatial distribution of age-related differences in structure-function coupling also recapitulated patterns of evolutionary cortical expansion. Age-related increases in coupling were observed primarily in highly expanded association cortex, while age-related decreases in coupling were observed in highly conserved sensory-motor cortex (*r*=0.39, *p_spin_*=0.002; **Fig. 3E**).

**Figure 3.**
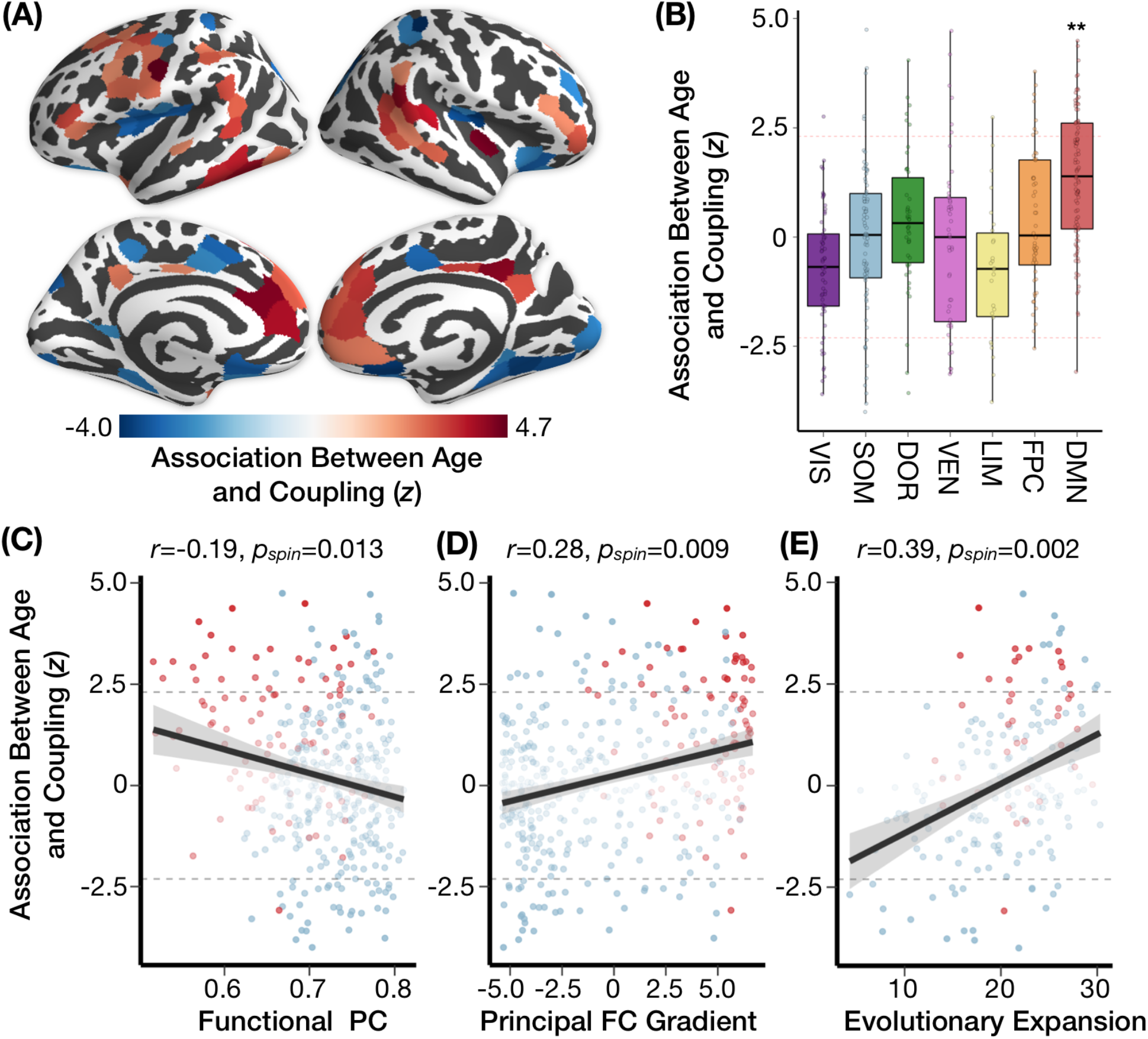
Hierarchy-dependent development of structure-function coupling. Age-related differences in structure-function coupling were broadly distributed across the cerebral cortex. (**A**) Age-related increases in structure-function coupling were observed bilaterally in the temporo-parietal junction and prefrontal cortex, while age-related decreases in coupling were observed in visual, motor and insular cortex. (**B**) Notably, age-related increases in coupling were disproportionately enriched within the default mode network compared to other functional systems (*F*=12.54, *p*<10^−10^). (**C**) The magnitude of age-related differences in structure-function coupling was significantly correlated with the functional participation coefficient (PC), (**D**) the functional gradient from unimodal to transmodal processing, and (**E**) evolutionary expansion of cortical surface area. The significance of regional correlations was evaluated using non-parametric spatial permutation testing (denoted *p_spin_*). Red points in C-E correspond to default mode regions, while blue points correspond to brain regions in other functional systems.

### Longitudinal increases in structure-function coupling are associated with changes in the regional diversity of functional connectivity

To determine whether age-related changes in structure-function coupling were reliably capturing intra-individual developmental change, we evaluated longitudinal changes in structure-function coupling using a sub-sample of participants who returned for follow-up approximately 1.7 years after baseline assessment (*n*=294). We observed a significant correspondence between cross-sectional and longitudinal age effects on structure-function coupling estimated with a linear mixed effects model (*r*=0.65, *p_spin_*<0.001; **Fig. 4A**).

**Figure 4.**
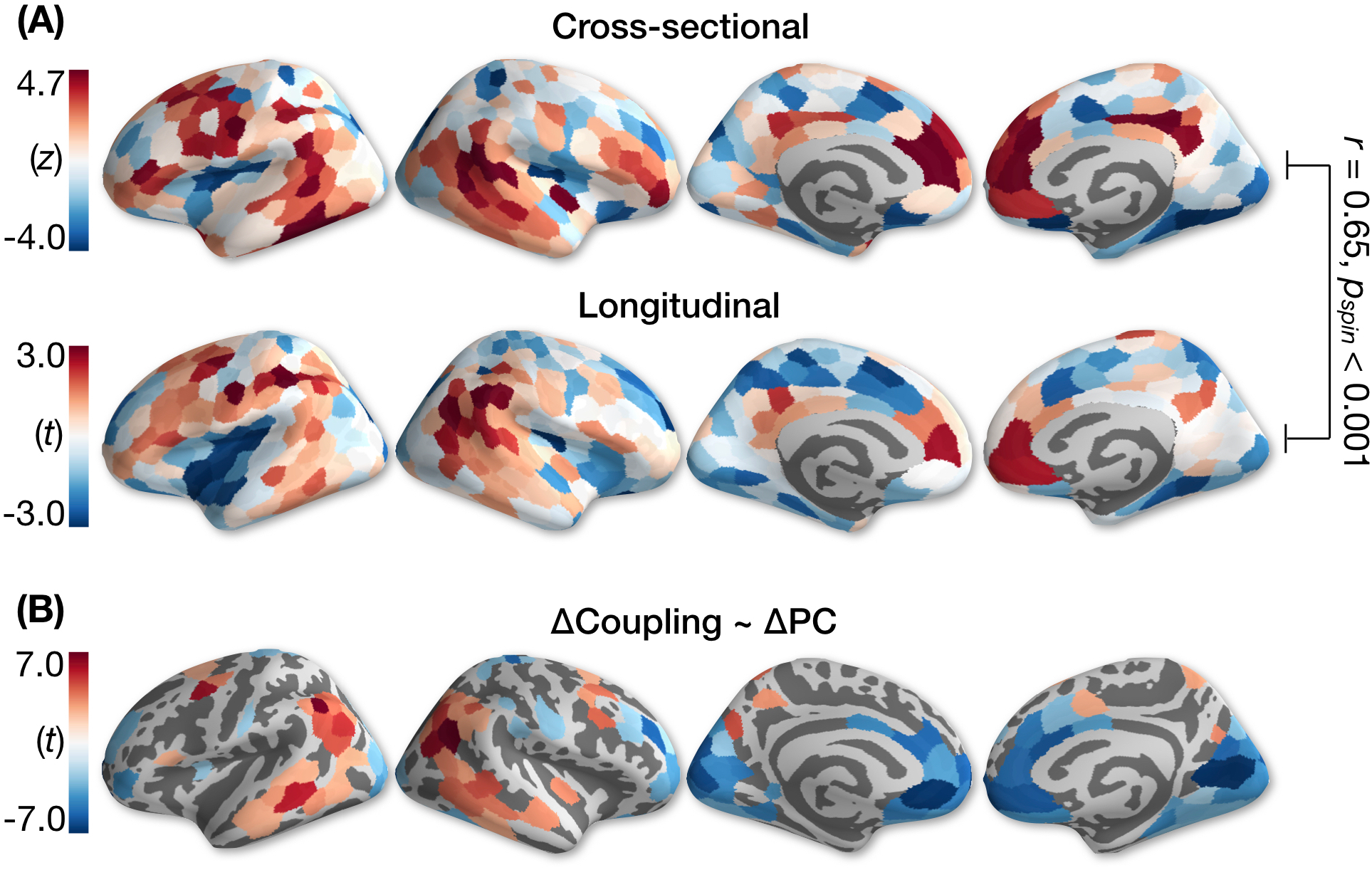
Longitudinal change in coupling is associated with longitudinal changes in the diversity of regional functional connectivity. (**A**) We observed a significant correspondence between cross-sectional (*n*=727) and longitudinal age effects on structure-function coupling estimated with a linear mixed effects model (*n*=294). (**B**) We used linear regression to test whether longitudinal change in coupling was associated with longitudinal change in the functional participation coefficient. In frontoparietal and lateral temporal regions, longitudinal increases in coupling were associated with higher participation coefficient. In medial occipital and medial prefrontal regions, longitudinal increases in structure-function coupling were associated with decreased participation coefficient.

Next, we evaluated how intra-individual development of structure-function coupling was associated with intra-individual changes in the diversity of regional connectivity. We focused on developmental changes in the participation coefficient because it captures how a brain region’s connections are distributed across functionally specialized sub-networks underlying perception, attention, and executive control. We used linear regression to test whether longitudinal change in coupling was associated with longitudinal change in the structural or functional participation coefficient. Notably, we found that longitudinal changes in structure-function coupling were associated with longitudinal changes in the functional participation coefficient in distributed higher-order association areas, including dorsomedial prefrontal, inferior parietal, and lateral temporal cortex (**Fig. 4B**). Specifically, longitudinal increases in coupling within dorsal prefrontal and inferior parietal regions were associated with increased inter-modular integration, while increased coupling in medial occipital and medial prefrontal cortex were associated with decreased inter-modular diversity (functional segregation). In contrast, only limited associations between longitudinal change in structure-function coupling and the structural participation coefficient were observed (*Supplementary Information*).

### Individual differences in structure-function coupling are associated with executive function

Next, we sought to understand the implications of individual differences in structure-function coupling for behavior. Specifically, we investigated whether structure-function coupling during a working memory task could explain executive performance measured on a computerized cognitive battery administered separately from the scanning session. While controlling for age, sex, and in-scanner head motion, we found that better executive performance was associated with higher structure-function coupling in the rostrolateral prefrontal cortex, posterior cingulate, and medial occipital cortex, and with lower structure-function coupling in somatosensory cortex (**Fig. 5A**). Regional associations between coupling and in-scanner performance on the *n*-back working memory task (*d*’) were highly consistent (**Supplementary Fig. 1**). Notably, the strength of this association between regional coupling and executive performance was significantly correlated with that region’s position along the functional hierarchy from unimodal to transmodal processing: higher structure-function coupling in transmodal regions of frontoparietal and default networks was associated with better performance on executive tasks (*r*=0.25, *p_spin_*=0.005). Furthermore, higher structure-function coupling in the right rostrolateral prefrontal cortex partially mediated age-related improvements in executive function (**Fig. 5B**; bootstrapped *p*=0.01). Regional associations between coupling and cognitive performance were specific to the executive domain: we observed no associations between coupling and social cognition, and stucture-function coupling was associated with memory performance in only four cortical regions. These results suggest that structure-function coupling in transmodal areas underpins individual differences in executive processes including working memory, attention and abstract reasoning.

**Figure 5.**
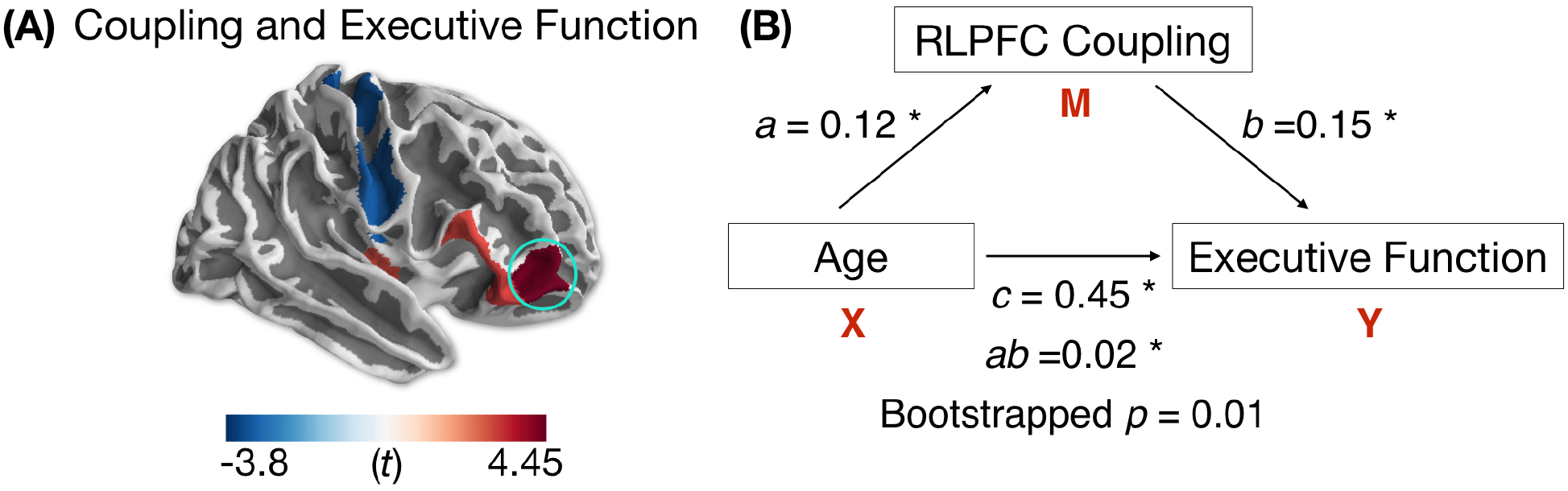
Individual differences in structure-function coupling are associated with executive performance. (**A**) We found that executive performance was associated with higher structure-function coupling in the rostrolateral prefrontal cortex, anterior insula, posterior cingulate, and medial occipital cortex, while better performance was associated with lower structure-function coupling in areas of somatomotor cortex. (**B**) Higher structure-function coupling in the right rostrolateral prefrontal cortex (RLPFC) partially mediated age-related improvements in executive function (circled, bootstrapped *p*=0.01). Mediation results are shown as standardized regression coefficients. Significance of the indirect effect (*ab*=0.02) was assessed using 95% bootstrapped confidence intervals [0.006-0.034].

### Sensitivity Analyses

As a final step, we performed sensitivity analyses to evaluate whether our results were robust to a number of methodological variations. Spatial variability and age-related changes in structure-function coupling were highly consistent across methodological approaches, including (*i*) using deterministic tractography and network communicability as a measure of structural connectivity strength that captures communication through indirect connections (**Supplementary Fig. 2**), (*ii*) extracting functional connectivity only from task blocks with high working memory load (1-back and 2-back) instead of the full task time-series (**Supplementary Fig. 3**), and (*iii*) accounting for inter-regional distance when quantifying structure-function coupling (**Supplementary Fig. 4**).

We also evaluated whether regional patterns of structure-function coupling showed a similar organization during the *n*-back working memory task and at rest. The spatial distribution of structure-function coupling was globally similar during *n*-back and rest when averaging across individuals (*r*=0.95, *p_spin_*<0.001). However, we observed greater intra-individual variability in regional coupling when assessing the correlation between *n*-back and resting-state coupling for each participant (mean *r*=0.53; **Supplementary Fig. 5**). Further, regional variability in structure-function coupling during *n*-back was more robustly associated with individual differences in executive performance compared to coupling during rest (*Supplementary Information*).

## DISCUSSION

We leveraged multimodal neuroimaging in a large sample of youth to characterize how structure-function coupling evolves in development and reflects macroscale cortical hierarchies. Consistent with previous work characterizing the targeted expansion and remodeling of transmodal cortex in both primate evolution and human development, we observed age-related differences in coupling localized within a unique subset of transmodal regions spanning higher-order association networks. These findings fill a critical gap in our understanding of how white matter architecture develops during human adolescence to support coordinated neural activity underlying executive processing.

Cortical hierarchy has provided a unifying principle for understanding the multiscale organization of primate cortical anatomy and function (2, 8, 23). Anatomical hierarchies of intracortical myelin (24) and laminar patterns of inter-areal projections (25) have been shown to align with hierarchies of functional (2) and transcriptional (24) specialization. Here, we provide evidence that these cortical hierarchies are in part determined by anatomical constraints on functional communication, whereby highly myelinated sensory areas exhibit strong structure-function coupling, and less myelinated association areas exhibit weak structure-function coupling. The convergence of structural and functional connectivity profiles in unimodal sensory regions suggests that functional communication is directly supported by local white matter pathways. In contrast, the divergence of structural and functional connectivity profiles in transmodal regions suggests that functional communication is untethered by structural constraints, relying more on polysynaptic (indirect) structural connections or circuit-level modulation of neural signals.

Lower structure-function coupling in transmodal brain regions may also support functional flexibility and dynamic recruitment during diverse task demands (26). One important exception to this trend was observed in transmodal regions of the default mode network, such as the medial prefrontal cortex, which exhibited both functionally segregated processing and relatively strong structure-function coupling. Tightly coupled structural and functional connectivity within transmodal regions of the medial prefrontal cortex could support efficient communication among strongly inter-connected association areas within the default mode network. Further, high structure-function coupling in local hubs of the default network could reduce competitive interference among central executive and task-negative networks (27), allowing for the suppression of internally-generated thoughts while maintaining and manipulating information in working memory.

Developmental changes in coupling were preferentially localized within transmodal areas of frontoparietal and default mode networks, recapitulating evolutionary patterns of cortical areal expansion. In addition to having expanded association cortex relative to other primates, humans exhibit slower axonal myelination in association cortex during childhood, characterized by a prolonged period of maturation that extends into early adulthood (5). As posited by the tethering hypothesis (18), this protracted development provides an extended window for the activity-dependent remodeling of distributed neural circuits in transmodal association cortex, which may be critical for the maturation of complex cognitive abilities in humans. In our study, longitudinal changes in structure-function coupling in transmodal cortex were associated with developmental increases in the diversity of inter-modular functional connectivity, underscoring the flexible and integrative role of these brain regions within the network.

One outstanding question concerns whether existing white matter architecture drives future changes in functional connectivity, or whether functional circuit changes sculpt the development of specific wiring patterns. We speculate that developmental changes in structure-function coupling could reflect processes of neural plasticity, such as the activity-dependent myelination of axons linking functionally coupled regions (28, 29). Alternatively, early myelination of axons could enhance signal conduction velocity and fidelity, enhancing neural signal-to-noise ratio (SNR) and the coordination of distributed neural activity (29). Longitudinal inferences in our study were limited by only two time-points of imaging data, precluding the characterization of lead-lag relationships between structural and functional brain connectivity. Future studies could leverage dense sampling of individuals during sensitive periods of development to delineate lead-lag relationships in the maturation of structural and functional connectivity within specialized circuits.

Our results also suggest that structure-function coupling has implications for individual differences in executive function. The rostrolateral prefrontal cortex (RLPFC) has been consistently linked with abstract reasoning (30) and the hierarchical control of goal-directed behavior (31). From childhood through early adulthood, the development of structural and functional connectivity between the RLPFC and lateral parietal cortex has been associated with improvements in abstract reasoning ability (30, 32). In this study, we extend these findings by showing that individual differences in RLPFC structure-function coupling partially mediate age-related improvements in executive functioning. The capacity of RLPFC to support executive processing may be understood through its role in integrating information between frontoparietal and dorsal attention networks to regulate perceptual attention (33).

Despite the strengths of this study, two potential limitations should be noted. First, accurately reconstructing the complexity of human white matter pathways from diffusion MRI and tractography remains challenging. Diffusion tractography algorithms face a well-characterized trade-off between connectome specificity and sensitivity (34). In this study, we attempted to overcome these limitations by replicating results with both deterministic and probabilistic tractography methods, while also applying a stringent consistency-based thresholding procedure to minimize the influence of false-positive connections (35). Second, motion artifact remains an important confound for all neuroimaging-based studies of brain development (36, 37). In addition to rigorous quality assurance protocols and extensively validated image processing designed to mitigate the influence of head motion on functional connectivity (38), we address this issue by quantifying and controlling for the influence of in-scanner head motion in all group-level analyses.

### Conclusion

By quantifying regional patterns of structure-function coupling and characterizing their development during adolescence, our results inform network-level mechanisms of plasticity that support cognitive maturation. Further, describing how underlying white matter architecture develops to support coordinated neural activity underlying executive function may offer critical insights into the basis for many sources of adolescent morbidity and mortality, such as risk-taking and diverse neuropsychiatric syndromes, which are prominently associated with failures of executive function.

## MATERIALS AND METHODS

Neuroimaging was conducted as part of the PNC (39). All participants included in this study were medically healthy, were not taking psychotropic medication at the time of study, and passed strict quality-assurance procedures for four imaging modalities including T1-weighted structural images, DWI, rs-fMRI, and *n*-back fMRI. The final sample included 727 youths ages 8–23 years old (420 females; mean=15.9, s.d.=3.2). From the original study sample, 147 typically developing youth returned for longitudinal neuroimaging assessments approximately 1.7 years after baseline (83 females; 294 total scans). For further details regarding image pre-processing and brain network construction, see *Supplementary Information*.

To evaluate the relationship between structure-function coupling and previously characterized cortical hierarchies, evolutionary cortical areal expansion (3) and the principal gradient of intrinsic functional connectivity (2) were extracted from publicly available atlases. The significance of the spatial correspondence between brain maps was estimated using a conservative spatial permutation test, which generates a null distribution of randomly rotated brain maps that preserve spatial covariance structure of the original data (21) (denoted *p_spin_*).

We used penalized splines within a generalized additive model (GAM) to estimate linear and nonlinear age-related changes in structure-function coupling for each brain region. Importantly, the GAM estimates nonlinearities using restricted maximum likelihood (REML), penalizing nonlinearity in order to avoid over-fitting the data (40). To evaluate regional associations between structure-function coupling and executive function, executive performance was measured as a factor score summarizing accuracy across mental flexibility, attention, working memory, verbal reasoning, and spatial ability tasks administered as part of the Penn Computerized Neurocognitive Battery (*Supplementary Information*).

Longitudinal intra-individual change in coupling and the participation coefficient were calculated as the difference in regional brain measures between timepoints. Baseline age, sex, mean relative frame-wise displacement, and the number of years between timepoints were included as additional co-variates in linear regression models.

## ACKNOWLEDGEMENTS

This study was supported by grants F31MH115709 (G.L.B.) and R01MH113550 (T.D.S. and D.S.B.) from the National Institute of Mental Health (NIMH). The PNC was supported by MH089983 and MH089924. Additional support was provided by R01MH107703 (T.D.S.), R01MH112847 (R.T.S. and T.D.S.), and R01MH107235 (R.C. G.), P50MH096891 (R.E.G.), K01MH102609 (D.R.R.), R01NS085211 (R.T.S.), RF1MH116920 (D.J.O., T.D.S. and D.S.B.), the Dowshen Program for Neuroscience, and the Lifespan Brain Institute at the Children’s Hospital of Philadelphia.

## REFERENCES

1. Huntenburg JM, Bazin P-L, Margulies DS (2018) Large-Scale Gradients in Human Cortical Organization. Trends in Cognitive Sciences 22(1):21–31.

2. Margulies DS, et al. (2016) Situating the default-mode network along a principal gradient of macroscale cortical organization. PNAS 113(44):12574–12579.

3. Hill J, et al. (2010) Similar patterns of cortical expansion during human development and evolution. PNAS 107(29):13135–13140.

4. Sotiras A, et al. (2017) Patterns of coordinated cortical remodeling during adolescence and their associations with functional specialization and evolutionary expansion. PNAS:201620928.

5. Miller DJ, et al. (2012) Prolonged myelination in human neocortical evolution. PNAS 109(41):16480–16485.

6. Petanjek Z, et al. (2011) Extraordinary neoteny of synaptic spines in the human prefrontal cortex. PNAS 108(32):13281–13286.

7. Larsen B, Luna B (2018) Adolescence as a neurobiological critical period for the development of higher-order cognition. Neuroscience & Biobehavioral Reviews 94:179–195.

8. Felleman DJ, Van Essen DC (1991) Distributed hierarchical processing in the primate cerebral cortex. Cereb Cortex 1(1):1–47.

9. Passingham RE, Stephan KE, Kötter R (2002) The anatomical basis of functional localization in the cortex. Nat Rev Neurosci 3(8):606–616.

10. Bassett DS, Sporns O (2017) Network neuroscience. Nat Neurosci 20(3):353–364.

11. Shen K, et al. (2012) Information processing architecture of functionally defined clusters in the macaque cortex. J Neurosci 32(48):17465–17476.

12. Saygin ZM, et al. (2012) Anatomical connectivity patterns predict face selectivity in the fusiform gyrus. Nat Neurosci 15(2):321–327.

13. Honey CJ, et al. (2009) Predicting human resting-state functional connectivity from structural connectivity. Proceedings of the National Academy of Sciences 106(6):2035–2040.

14. Mišić B, et al. (2016) Network-Level Structure-Function Relationships in Human Neocortex. Cereb Cortex 26(7):3285–3296.

15. Goñi J, et al. (2014) Resting-brain functional connectivity predicted by analytic measures of network communication. Proceedings of the National Academy of Sciences 111(2):833–838.

16. Di Martino A, et al. (2014) Unraveling the Miswired Connectome: A Developmental Perspective. Neuron 83(6):1335–1353.

17. Stephan KE, Friston KJ, Frith CD (2009) Dysconnection in Schizophrenia: From Abnormal Synaptic Plasticity to Failures of Self-monitoring. Schizophr Bull 35(3):509–527.

18. Buckner RL, Krienen FM (2013) The evolution of distributed association networks in the human brain. Trends in Cognitive Sciences 17(12):648–665.

19. Greene AS, Gao S, Scheinost D, Constable RT (2018) Task-induced brain state manipulation improves prediction of individual traits. Nature Communications 9(1):2807.

20. Schaefer A, et al. (2018) Local-Global Parcellation of the Human Cerebral Cortex from Intrinsic Functional Connectivity MRI. Cereb Cortex 28(9):3095–3114.

21. Alexander-Bloch AF, et al. (2018) On testing for spatial correspondence between maps of human brain structure and function. NeuroImage 178:540–551.

22. Guimerà R, Amaral LAN (2005) Cartography of complex networks: modules and universal roles. J Stat Mech 2005(P02001):nihpa35573.

23. Markov NT, et al. (2014) Anatomy of hierarchy: feedforward and feedback pathways in macaque visual cortex. J Comp Neurol 522(1):225–259.

24. Burt JB, et al. (2018) Hierarchy of transcriptomic specialization across human cortex captured by structural neuroimaging topography. Nat Neurosci 21(9):1251–1259.

25. Barbas H, Rempel-Clower N (1997) Cortical structure predicts the pattern of corticocortical connections. Cereb Cortex 7(7):635–646.

26. Yeo BTT, et al. (2015) Functional Specialization and Flexibility in Human Association Cortex. Cereb Cortex 25(10):3654–3672.

27. Hampson M, Driesen N, Roth JK, Gore JC, Constable RT (2010) Functional connectivity between task-positive and task-negative brain areas and its relation to working memory performance. Magn Reson Imaging 28(8):1051–7.

28. Gibson EM, et al. (2014) Neuronal Activity Promotes Oligodendrogenesis and Adaptive Myelination in the Mammalian Brain. Science 344(6183):1252304.

29. Mount CW, Monje M (2017) Wrapped to Adapt: Experience-Dependent Myelination. Neuron 95(4):743–756.

30. Wendelken C, Ferrer E, Whitaker KJ, Bunge SA (2016) Fronto-Parietal Network Reconfiguration Supports the Development of Reasoning Ability. Cereb Cortex 26(5):2178–2190.

31. Desrochers TM, Chatham CH, Badre D (2015) The necessity of rostrolateral prefrontal cortex for higher-level sequential behavior. Neuron 87(6):1357–1368.

32. Wendelken C, et al. (2017) Frontoparietal Structural Connectivity in Childhood Predicts Development of Functional Connectivity and Reasoning Ability: A Large-Scale Longitudinal Investigation. J Neurosci 37(35):8549–8558.

33. Dixon ML, et al. (2018) Heterogeneity within the frontoparietal control network and its relationship to the default and dorsal attention networks. PNAS 115(7):E1598–E1607.

34. Zalesky A, et al. (2016) Connectome sensitivity or specificity: which is more important? NeuroImage 142:407–420.

35. Roberts JA, Perry A, Roberts G, Mitchell PB, Breakspear M (2017) Consistency-based thresholding of the human connectome. NeuroImage 145:118–129.

36. Satterthwaite TD, et al. (2013) Heterogeneous impact of motion on fundamental patterns of developmental changes in functional connectivity during youth. Neuroimage 83:45–57.

37. Baum GL, et al. (2018) The impact of in-scanner head motion on structural connectivity derived from diffusion MRI. NeuroImage 173:275–286.

38. Ciric R, et al. (2018) Mitigating head motion artifact in functional connectivity MRI. Nat Protoc 13(12):2801–2826.

39. Satterthwaite TD, et al. (2014) Neuroimaging of the Philadelphia neurodevelopmental cohort. Neuroimage 86:544–553.

40. Wood SN (2011) Fast stable restricted maximum likelihood and marginal likelihood estimation of semiparametric generalized linear models. Journal of the Royal Statistical Society: Series B (Statistical Methodology) 73(1):3–36.

