## Supplemental Information for "Development of structure-function coupling in human brain networks during youth"

### SUPPLEMENTARY INFORMATION

*Participants*

Neuroimaging and behavioral data were originally obtained from 1,601 youth who participated in the Philadelphia Neurodevelopmental Cohort (PNC), a large community-based study of brain development (1, 2). From this original sample, 340 participants were excluded based on health criteria, including psychoactive medication use at the time of study, medical problems that could impact brain function, a history of psychiatric hospitalization, and gross structural brain abnormalities (3, 4). Of the remaining 1,261 participants, 45 were excluded for poor T1-weighted image quality, which assessed with both manual and automated quality assurance procedures. Of the remaining 1,216 participants, 267 were excluded for poor quality or missing *n*-back functional connectivity data. From the remaining sample, 149 participants were excluded for poor quality or missing resting-state functional connectivity data, and 160 were then removed for poor quality or missing diffusion-weighted imaging data. After these exclusions, 740 participants remained in the study sample. Lastly, 3 participants were excluded due to poor atlas coverage in native diffusion space, and 10 more participants were excluded for having fully disconnected nodes in structural brain networks. In sum, following rigorous quality assurance procedures for structural imaging, diffusion imaging, n-back task fMRI, and resting-state fMRI, we retained 727 participants in the study sample between the ages of 8 and 23 years old (mean= 15.9 years, s.d. = 3.2, 420 females). See *MRI quality assurance* section below for further details regarding subject exclusion criteria for each imaging modality.

*Cognitive Assessment*

The Penn computerized neurocognitive battery (Penn CNB) was administered to all participants. The CNB consists of 14 tests adapted from tasks applied in functional neuroimaging to evaluate a broad range of cognitive domains (5). These domains include executive control (abstraction and flexibility, attention, working memory), complex cognition (verbal reasoning, nonverbal reasoning, spatial processing), episodic memory (verbal, facial, spatial), social cognition (emotion identification, emotion intensity differentiation, age differentiation), and motor speed. Accuracy and speed for each test were z-transformed. Cognitive performance was summarized by a recent factor analysis of CNB data (6), which delineated three factors corresponding to the accuracy of executive function, episodic memory, and social cognition. We evaluated associations between executive accuracy and structure-function coupling. Additional sensitivity analyses were performed with social cognition and memory accuracy scores (*Supplemental Results*).

*Image Acquisition*

All MRI scans were acquired on the same 3T Siemens Tim Trio whole-body scanner and 32-channel head coil at the Hospital of the University of Pennsylvania. Prior to DWI acquisition, a 5-minute magnetization-prepared, rapid acquisition gradient-echo T1-weighted (MPRAGE) image (TR 1810 ms, TE 3.51 ms, FOV 180 × 240 mm, matrix 256 × 192, effective voxel resolution of 1 × 1 × 1 mm) was acquired (1). DWI scans were acquired using a twice- refocused spin-echo (TRSE) single-shot echoplanar imaging (EPI) sequence (TR = 8100ms, TE = 82ms, FOV = 240mm$^2$ / 240mm$^2$; Matrix = RL: 128, AP:128, Slices:70, in-plane resolution (x and y) 1.875 mm$^2$; slice thickness = 2mm, gap = 0; flip angle = 90°/180°/180°, volumes = 71, GRAPPA factor = 3, bandwidth = 2170 Hz/pixel, PE direction = AP). This sequence used a four-lobed diffusion encoding gradient scheme combined with a 90-180-180 spin-echo sequence designed to minimize eddy-current artifacts (1). For DWI acquisition, a 64-direction set was divided into two independent 32-direction imaging runs in order make scan duration more tolerable for young subjects. Two consecutive 32-direction acquisitions were merged into a single 64-direction time-series. The complete sequence consisted of 64 diffusion-weighted directions with b = 1000s/ mm$^2$ and 7 interspersed scans where b = 0 s/ mm$^2$. The total duration of DWI scans was approximately 11 minutes. The imaging volume was prescribed in axial orientation covering the entire cerebrum with the topmost slice just superior to the apex of the brain (1).. A map of the main magnetic field (i.e., B0) was derived from a double-echo, gradient-recalled echo (GRE) sequence, allowing us to estimate field distortions in each dataset (1). All subjects also completed blood oxygen level-dependent (BOLD-weighted) *n*-back task fMRI (12 minute duration) and resting-state fMRI (6 minute duration) with identical acquisition parameters (TR=3000 ms; TE=32 ms; flip angle=90°; FOV=192 × 192 mm; matrix = 64×64; slices=46; slice thickness=3 mm; slice gap=0 mm; effective voxel resolution=3.0 × 3.0 × 3.0mm) (1).

*MRI Quality Assurance*

Images from each modality underwent a rigorous quality assurance procedure. Each subject's T1-weighted anatomical image quality was independently rated by three highly trained image analysts (7). Image quality ratings were averaged across the three raters as a summary measure of image quality. After processing T1-weighted images with the ANTS Cortical Thickness Pipeline (8), regional cortical thickness outliers were identified as 2.5 SD above or below mean regional values. This automated quality assurance procedure flagged additional structural images, which were subsequently inspected manually by trained specialists to determine whether images were usable or not.

Prior to diffusion image processing, all raw DWI datasets were subject to a rigorous manual quality assessment procedure involving visual inspection of all 71 volumes (9). Each volume was evaluated for the presence of artifact, with the total number of volumes impacted summed over the series. This scoring was based on previous work describing the impact of removing image volumes when estimating the diffusion tensor (10, 11). Data was considered ‘‘Poor’’ if more than 14 (20%) volumes contained artifact, ‘‘Good’’ if it contained 1-14 volumes with artifact, and ‘‘Excellent’’ if no visible artifacts were detected in any volumes. All subjects included in the present study had diffusion datasets identified as ‘‘Good’’ or ‘‘Excellent” (9). As described below, even after this rigorous quality assurance, mean relative displacement between interspersed b = 0 volumes was as included as a nuisance covariate in all group-level analyses.

Participants were also excluded for poor fMRI data if maximum relative root-mean-square framewise displacement exceeded 6mm, or if the mean relative root-mean-square framewise displacement exceeded 0.5mm during *n*-back or resting-state scans (12).

*Structural image processing*

A study-specific template was generated from a sample of 120 PNC subjects balanced across sex, race, and age bins using the *buildtemplateparallel* procedure in ANTS (13). All images that did not pass manual inspection were removed from the analysis. Each subject's high-resolution structural image was processed using the ANTS Cortical Thickness Pipeline (8). Following bias field correction (14), each structural image was diffeomorphically registered to the study-specific PNC template using the top-performing SYN deformation provided by ANTS (15). Study-specific tissue priors were used to guide brain extraction and segmentation of the subject's structural image (16, 17).

*Diffusion image processing*

A mask in subject diffusion space was defined by registering a binary mask of a standard fractional anisotropy (FA) map (FMRIB58 FA) to each subject's diffusion reference image (mean b=0) using FLIRT (18). This mask was provided as input to FSL eddy in addition to the non-brain extracted dMRI image. Eddy currents and subject motion were estimated and corrected using the FSL eddy tool (version 5.0.5) (19). This procedure uses a Gaussian Process to simultaneously model the effects of eddy currents and head motion on diffusion-weighted volumes, resampling the data only once. Diffusion gradient vectors were also rotated to adjust for subject motion estimated by eddy (20). After the field map was estimated, distortion correction then was applied to dMRI images using FSL's FUGUE (21).

*Diffusion model fitting, probabilistic tractography, and structural brain network construction*

A ball-and-sticks diffusion model was fitted to each subject's DWI data using FSL bedpostx, which uses Markov chain Monte Carlo sampling to build distributions on principal fiber orientation and diffusion parameters at each voxel (22, 23). This allowed us to model up to two crossing fibers per voxel, enhancing sensitivity to more complex white matter architecture. Probabilistic tractography was run using FSL probtrackx, which repetitively samples voxel-wise fiber orientation and diffusion parameter distributions to model the spatial trajectory and strength of white matter connectivity between specified seed and target regions (23).

Seed and target regions were defined in native diffusion space after co-registering a standard 400-region brain parcellation of cortical gray matter (24) to the PNC study-specific template and the high-resolution T1-weighted image. White matter and cerebro-spinal fluid segmentations defined using the ANTS Cortical Thickness pipeline were also co-registered to native diffusion space to serve as waypoint and exclusion masks, respectively. The white matter boundary was defined by using the *fslmaths -edge* function on the ANTS white matter segmentation, and this white matter edges edge image was dilated by 1 voxel (1.875 mm) to generate a ribbon along the gray-white boundary in native diffusion space. Seed and target regions were defined by masking the original gray matter ROIs by the dilated WM edge (25, 26). A termination mask of superficial gray matter was also generated by subtracting the original gray matter ROIs from the gray-white boundary ROIs.

Each cortical region defined along the gray-white boundary was selected as a seed region, and its connectivity strength to each of the other 399 regions was calculated using probabilistic tractography. At each seed voxel, 1000 samples were initiated (25–27). Default tracking parameters were applied otherwise (a step-length of 0.5mm, 2000 steps maximum, curvature threshold of 0.02). To increase the biological plausibility of white matter pathways reconstructed with probabilistic tractography, streamlines were terminated if they entered superficial gray matter, and discarded if they traversed cerebro-spinal fluid (CSF) in ventricles or re-entered the seed region (26). This fiber tracking procedure allowed us to construct a weighted n × n connectivity matrix for each participant (see **Fig. 1**), where connection weights were defined as the number of probabilistic streamlines connecting each pair of brain regions (25, 26). Edge weights in each subject's connectivity matrix were normalized by the total weight of network connections in order to delineate intrinsic topological differences between subjects (25–28).

*Consistency-based thresholding of structural connectivity matrices*

Probabilistic tractography yields dense weighted networks that contain a large number of potentially spurious connections. Several approaches exist for mitigating the influence of false positive and false negative connections reconstructed in structural connectomes. While one common thresholding approach involves removing a subset of the weakest edges in a group-average connection matrix (29), this approach often results in the elimination of relatively weak, long-range connections that may play an important role in brain network topology (30, 31). In contrast, consistency-based thresholding considers the coefficient of variation (CV) for each network connection in the study sample, and retains both short- and long-range connections that are consistently reconstructed across subjects (31). In this study, each subject's structural connectivity matrix was thresholded at the 75$^{th}$ percentile for edge consistency, pruning the most inconsistent connections identified in the top quartile for CV.

*Sensitivity analysis using deterministic tractography*

To ensure that our results were not influenced by the relatively high rate of reconstructing false-positive connections using probabilistic fiber tracking methods (32), we also generated structural brain networks using deterministic tractography. Whole-brain deterministic fiber tracking was implemented for each participant in DSI Studio (33) using a modified fiber assessment by continuous tracking (FACT) algorithm with Euler interpolation, initiating 1,000,000 streamlines after removing all streamlines with length less than 10mm or greater than 400mm. Fiber tracking was performed with an angular threshold of 45°, a step size of 0.9375mm, and a fractional anisotropy (FA) threshold determined empirically by Otzu's method, which optimizes the contrast between foreground and background (33). Edge weights were initially defined using number of deterministic streamlines connecting any pair of nodes (25, 26, 34). Deterministic tractography yielded relatively sparse brain networks, which was problematic for calculating regional structure-function coupling profiles due to a low number of non-zero edges in each regional connectivity profile. Further, forty-one participants were excluded from analysis due to having at least one fully disconnected node in their structural brain network, precluding estimation of structure-function coupling. In order to evaluate spatial variation and age-related changes in structure-function coupling in the remaining 686 participants, we calculated the communicability for each network connection, which captures the communication capacity through both direct and indirect connectivity between each pair of brain regions (35). This results in a fully-connected communicability matrix, where edge weights reflect the weighted sum of both direct and indirect pathways between regions, where shorter paths with stronger connections are weighted more heavily. To enhance biological plausibility of structural brain networks, we applied the same consistency-based threshold used for networks derived from probabilistic tractography, yielding an average network density of 54.3% (SD=5.8).

*Fractal n-back fMRI task*

Performance of the fractal *n*-back working memory task reliably activates the frontoparietal executive system (36). Furthermore, a fractal version of the *n*-back task is particularly useful for delineating the development of working memory without the confound of lexical processing (37–39). Each task condition included a series of 60 fractal stimuli separated over three 20-stimulus blocks. Each stimulus was presented for 500ms with inter-stimulus intervals of 2500ms (total of 60s per block). The three task conditions, ordered according to increasing working memory load, were the 0-back, 1-back, and 2-back conditions. During the 0-back condition, participants were instructed to press a button in response to a single target stimulus. During the 1-back condition, subjects were instructed to press a button if the current stimulus matched the previous stimulus. During the 2-back condition, subjects were instructed to press a button if the current stimulus matched the stimulus presented two trials prior. The overall ratio of target stimuli to foil stimuli was maintained over all conditions as 1:3, with a total 15 target stimuli and 45 foil stimuli in each condition. Prior to the scan session, a mock scanning session was conducted to acclimate subjects to the scan environment (1).

*fMRI processing*

Both *n*-back and resting-state functional images were processed using one of the top-performing pipelines for removal of motion-related artifact (40) within the XCP engine (41). Preprocessing steps included (a) correction for distortions induced by magnetic field inhomogeneities using FSL's FUGUE utility, (b) removal of the 4 initial volumes of each acquisition, (c) realignment of all volumes to a selected reference volume using MCFLIRT (18), (d) removal of and interpolation over intensity outliers in each voxel's time series using AFNI's 3DDESPIKE utility, (e) demeaning and removal of any linear or quadratic trends, and (f) co-registration of functional data to the high-resolution structural image using boundary-based registration (42). The artefactual variance in the data was modelled using a total of 36 parameters, including the six frame-wise estimates of motion, the mean signal extracted from eroded white matter and cerebrospinal fluid compartments, the mean signal extracted from the entire brain, the derivatives of each of these nine parameters, and quadratic terms of each of the nine parameters and their derivatives. Both the BOLD-weighted time series and the artefactual model time series were temporally filtered using a first-order Butterworth filter with a passband between 0.01 and 0.08 Hz (43).

*Functional connectome construction*

Following de-noising, functional connectivity between each pair of brain regions was quantified as the Pearson correlation coefficient between the mean regional BOLD time series. For each participant, an 400 × 400 weighted adjacency matrix encoding the connectome was constructed (see **Fig. 1**). Each node was assigned to one of seven canonical functional brain modules or communities defined by Yeo et al. (24, 44).

*Measuring structure-function coupling in human brain networks*

Regional connectivity profiles were extracted from each column of a participant's structural or functional connectivity matrix, and were represented as vectors of connectivity strength from a single network node to all other nodes in the network. Structure-function coupling was then measured as the Spearman rank correlation between nonzero elements of regional structural and functional connectivity profiles (45, 46). Regional indices of structure-function coupling were averaged across participants to create a mean regional coupling map (**Fig. 2A**).

*Evolutionary areal expansion and principal rs-FC gradient maps*

Evolutionary cortical surface area expansion between macaques and humans was estimated by measuring the surface deformation that would bring human cortical areas into spatial alignment with their macaque homologues, and extracted from a publically available atlas (47). The principal gradient of intrinsic functional connectivity, which reflects a functional hierarchy from unimodal sensory cortex to transmodal association cortex, was also extracted from a publicly available atlas (48).

*Spatial Permutation testing*

The significance of the spatial correspondence between structure-function coupling and other cortical properties (such as evolutionary expansion and rs-FC gradients) brain maps was estimated using a highly conservative spatial permutation test, which generates a null distribution of randomly rotated brain maps that preserve spatial covariance structure of the original data (49) Specifically, the mean structure-function coupling map (see **Fig. 2A**) was projected to an fsaverage6 spherical cortical surface and rotated randomly 1000 times, generating a distribution of “null” maps that preserve spatial neighborhood information. Structure-function coupling was extracted for each region in randomly rotated maps, and the Pearson correlation coefficient between regional coupling and other cortical measures (e.g., functional participation coefficient) was calculated to build a null distribution. The permutation-based *p*-value was calculated as the proportion of times that null correlation coefficients were greater than empirical correlation coefficients between regional measures (49). This spin test procedure was repeated using the map of age-related changes in structure-function coupling to determine whether age effects were significantly correlated with functional diversity, rs-FC hierarchy, and evolutionary areal expansion.

*Group-level statistical analysis*

We used penalized splines within a generalized additive model (GAM) to estimate linear and nonlinear age-related changes in structure-function coupling for each brain region. Importantly, the GAM estimates nonlinearities using restricted maximum likelihood (REML), penalizing nonlinearity in order to avoid over-fitting the data (50, 51). Within this model, we included covariates for sex and head motion during both diffusion and n-back scans. We controlled for multiple comparisons using the False Discovery Rate ($Q$<0.05).

*Longitudinal group-level analysis*

To determine whether age-related changes in structure-function coupling were reliably capturing within-subject developmental change, we evaluated longitudinal changes in structure-function coupling using a sub-sample of participants who returned for follow-up approximately 1.7 years after baseline assessment ($n$=294). Longitudinal developmental changes in structure-function were estimated using a linear mixed effects model (*nlme* package), including a random subject intercept term to account for repeated measurements.

We evaluated whether within-subject change in structure-function coupling was associated with the refinement of regional functional or structural connectivity profiles. Specifically, we tested a linear regression model with longitudinal change in coupling as the dependent variable, and longitudinal change in the structural or functional participation coefficient as dependent variables. Baseline age, sex, mean relative frame-wise displacement, and the number of years between time-points were included as additional co-variates in regression models. Longitudinal within-subject change in coupling and the participation coefficient were calculated as the difference in regional brain measures between baseline and follow-up assessments. Baseline age, sex, mean relative frame-wise displacement, and the number of years between time-points were included as additional co-variates in regression models. Results remained highly consistent when using residual change scores, or normalizing raw change scores within-subjects for regression testing.

*Mediation analysis*

Mediation analyses investigated whether age-related improvement in executive function was mediated by regional patterns of structure-function coupling. First, we regressed out the effects of nuisance covariates (sex and head motion) on the independent (*X*), dependent (*Y*), and mediating (*M*) variables. The normalized residuals were then used in our mediation analysis. The significance of the indirect effect was evaluated using bootstrapped confidence intervals within the R package *lavaan*.

Specifically, we examined the total effect of age on executive performance, the relationship between age and structure-function coupling (*a* path), the relationship between structure-function coupling and executive performance (*b* path), and the direct effect of age on executive performance after including structure-function coupling as a mediator in the model (*c′* path). The significance of the indirect effect (*ab*) of age on executive function through the proposed mediator (structure-function coupling) was tested using bootstrapping procedures, which minimize assumptions about the sampling distribution (52). This approach involves calculating indirect effects for each of 10,000 bootstrapped samples and then calculating the 95% confidence interval.

### SUPPELEMENTARY RESULTS

*Longitudinal development of structure-function coupling is associated with changes in the functional participation coefficient*

We used linear regression to test whether longitudinal change in coupling was associated with longitudinal change in the structural or functional participation coefficient. As previously noted, we found that longitudinal changes in structure-function coupling were associated with longitudinal changes in the functional participation coefficient in 151 distributed brain regions, including dorsomedial prefrontal, inferior parietal, and lateral temporal cortex (**Fig. 4B**). In contrast, only limited associations between longitudinal change in structure-function coupling and the structural participation coefficient were observed. Specifically, we found a negative association between change in coupling and change in the structural participation coefficient for 11 brain regions spanning bilateral visual, somatomotor, and medial prefrontal cortex. These results suggest that longitudinal increases in structure-function coupling were associated with decreased diversity of regional structural connectivity (increased segregation) in these brain regions.

*Sensitivity analyses*

We evaluated whether regional associations between structure-function coupling and executive performance were consistent when using in-scanner performance on the *n*-back fMRI task (*d′*) instead of performance on executive tasks administered separately with the Penn Computerized Neurocognitive Battery. *N*-back performance was assessed using *d′*, a composite measure that takes into account both correct responses and false positives to separate performance from response bias (36). Regional associations between structure-function coupling and *n*-back performance (*d′*) were highly correlated with associations between coupling and executive performance on a computerized battery (**Supplementary Fig. S1**, $r$=0.80, $p_{spin}$<0.001). These results suggest that structure-function coupling in transmodal areas underpins individual differences in executive processes including working memory, attention and abstract reasoning, and that these associations are not driven by epiphenomena of task fMRI.

Next, we conducted a thorough set of analyses to examine whether our results were dependent on specific methodological choices. First, due to the well-documented trade-off in connectome sensitivity and specificity with different fiber tracking methods, we evaluated whether our results were consistent when using deterministic tractography. Specifically, we calculated structure-function coupling using each participant's thresholded communicability matrix derived from deterministic tractography. We found a strong spatial correlation between mean regional structure-function coupling calculated using deterministic and probabilistic tractography methods for structural brain network construction (Pearson $r$=0.79, $p_{spin}$<0.001**; Supplementary Fig. 2A**). Age-related changes in structure-function coupling also remained highly consistent (**Supplementary Fig 2B**), and were distributed across superior temporal, parietal, cingulate, and prefrontal areas. Consistent with our main findings, we observed hierarchy-dependent development of structure-function coupling ($r$=0.34, $p_{spin}$=0.002; **Supplementary Fig. 2C**). Specifically, age-related increases in coupling were localized within transmodal areas of fronto-parietal and default networks, while age-related decreases in coupling were localized primarily within unimodal sensory areas.

Second, to ensure that our results were driven by working-memory related processing, we evaluated whether our results were consistent when measuring functional connectivity from BOLD time-series extracted only from task blocks with high working memory load (1-back and 2-back) instead of the full task time-series. Structure-function coupling was then quantified using this working memory-related FC and structural connectivity derived from probabilistic tractography. Mean regional structure-function coupling calculated with high working memory load was highly correlated with coupling calculated using the full *n*-back time-series ($r$=0.99, $p_{spin}$<0.001; **Supplementary Fig. 3A**). Age-related changes in structure-function coupling also remained highly consistent, and were distributed across superior temporal, parietal, cingulate, and prefrontal cortex (**Supplementary Fig 3B**). We observed hierarchy-dependent development of structure-function coupling $r$=0.25, $p_{spin}$=0.015; **Supplementary Fig. 3C**), with age-related increases in coupling localized within transmodal areas of fronto-parietal and default networks, and age-related decreases in coupling localized primarily within unimodal sensory areas.

Third, we evaluated regional structure-function coupling while accounting for the influence of inter-regional connection distance, which imposes well-characterized constraints on brain connectivity (53). We found a significant spatial correlation between mean structure-function coupling maps that did and did not account for inter-regional connection distance ($r$=0.47, $p_{spin}$<0.001; **Supplementary Fig. 4A**). When accounting for inter-regional distance, one notable difference was that transmodal regions in frontoparietal and default networks exhibited higher structure-function coupling, while unimodal sensory regions had relatively lower structure-function coupling. Despite these differences, age-related changes in structure-function coupling still remained highly consistent when accounting for inter-regional distance, and were distributed primarily in parietal, cingulate, and prefrontal cortex (**Supplementary Fig. 4B**). Despite subtle differences in the spatial organization of structure-function coupling when accounting for inter-regional connection distance, we still observed hierarchy-dependent development of structure-function coupling ($r$=0.40, $p_{spin}$<0.001; **Supplementary Fig. 4C**).

### Supplementary References


1. Satterthwaite TD, et al. (2014) Neuroimaging of the Philadelphia neurodevelopmental cohort. *Neuroimage* 86:544–553.

2. Satterthwaite TD, et al. (2016) The Philadelphia Neurodevelopmental Cohort: A publicly available resource for the study of normal and abnormal brain development in youth. *Neuroimage* 124(Pt B):1115–1119.

3. Merikangas KR, et al. (2010) Prevalence and treatment of mental disorders among US children in the 2001-2004 NHANES. *Pediatrics* 125(1):75–81.

4. Gur RE, et al. (2013) Incidental findings in youths volunteering for brain MRI research. *AJNR Am J Neuroradiol* 34(10):2021–2025.

5. Gur RC, et al. (2012) Age group and sex differences in performance on a computerized neurocognitive battery in children age 8-21. *Neuropsychology* 26(2):251–65.

6. Moore TM, Reise SP, Gur RE, Hakonarson H, Gur RC (2015) Psychometric Properties of the Penn Computerized Neurocognitive Battery. *Neuropsychology* 29(2):235–246.

7. Rosen AFG, et al. (2018) Quantitative assessment of structural image quality. *NeuroImage* 169:407–418.

8. Tustison NJ, et al. (2014) Large-scale evaluation of ANTs and FreeSurfer cortical thickness measurements. *NeuroImage* 99(Supplement C):166–179.

9. Roalf DR, et al. (2016) The impact of quality assurance assessment on diffusion tensor imaging outcomes in a large-scale population-based cohort. *NeuroImage* 125:903–919.

10. Chen Y, Tymofiyeva O, Hess CP, Xu D (2015) Effects of rejecting diffusion directions on tensor-derived parameters. *NeuroImage* 109:160–170.

11. Jones DK, Basser PJ (2004) Squashing peanuts and smashing pumpkins": How noise distorts diffusion-weighted MR data. *Magnetic Resonance in Medicine* 52(5):979–993.

12. Xia CH, et al. (2018) Linked dimensions of psychopathology and connectivity in functional brain networks. *Nature Communications* 9(1):3003.

13. Avants BB, et al. (2011) A reproducible evaluation of ANTs similarity metric performance in brain image registration. *Neuroimage* 54(3):2033–2044.

14. Tustison NJ, et al. (2010) N4ITK: improved N3 bias correction. *IEEE Trans Med Imaging* 29(6):1310–1320.

15. Klein A, et al. (2009) Evaluation of 14 nonlinear deformation algorithms applied to human brain MRI registration. *Neuroimage* 46(3):786–802.

16. Avants BB, Tustison NJ, Wu J, Cook PA, Gee JC (2011) An open source multivariate framework for n-tissue segmentation with evaluation on public data. *Neuroinformatics* 9(4):381–400.

17. Wang H, et al. (2013) Multi-Atlas Segmentation with Joint Label Fusion. *IEEE Trans Pattern Anal Mach Intell* 35(3):611–623.

18. Jenkinson M, Bannister P, Brady M, Smith S (2002) Improved optimization for the robust and accurate linear registration and motion correction of brain images. *Neuroimage* 17(2):825–841.

19. Andersson JLR, Sotiropoulos SN (2016) An integrated approach to correction for off-resonance effects and subject movement in diffusion MR imaging. *Neuroimage* 125:1063–78.

20. Leemans A, Jones DK (2009) The B-matrix must be rotated when correcting for subject motion in DTI data. *Magn Reson Med* 61(6):1336–1349.

21. Jenkinson M, Beckmann CF, Behrens TE, Woolrich MW, Smith SM (2012) Fsl. *Neuroimage* 62(2):782–790.

22. Behrens TEJ, et al. (2003) Characterization and propagation of uncertainty in diffusion-weighted MR imaging. *Magn Reson Med* 50(5):1077–88.

23. Behrens TEJ, Berg HJ, Jbabdi S, Rushworth MFS, Woolrich MW (2007) Probabilistic diffusion tractography with multiple fibre orientations: What can we gain? *NeuroImage* 34(1):144–155.

24. Schaefer A, et al. (2018) Local-Global Parcellation of the Human Cerebral Cortex from Intrinsic Functional Connectivity MRI. *Cereb Cortex* 28(9):3095–3114.

25. Baum GL, et al. (2017) Modular Segregation of Structural Brain Networks Supports the Development of Executive Function in Youth. *Current Biology* 27(11):1561-1572.e8.

26. Baum GL, et al. (2018) The impact of in-scanner head motion on structural connectivity derived from diffusion MRI. *NeuroImage* 173:275–286.

27. Li L, Rilling JK, Preuss TM, Glasser MF, Hu X (2012) The effects of connection reconstruction method on the interregional connectivity of brain networks via diffusion tractography. *Hum Brain Mapp* 33(8):1894–1913.

28. Gong G, et al. (2009) Age- and gender-related differences in the cortical anatomical network. *J Neurosci* 29(50):15684–93.

29. Rubinov M, Sporns O (2010) Complex network measures of brain connectivity: uses and interpretations. *Neuroimage* 52(3):1059–1069.

30. Drakesmith M, et al. (2015) Overcoming the effects of false positives and threshold bias in graph theoretical analyses of neuroimaging data. *NeuroImage* 118:313–333.

31. Roberts JA, Perry A, Roberts G, Mitchell PB, Breakspear M (2017) Consistency-based thresholding of the human connectome. *NeuroImage* 145:118–129.

32. Maier-Hein KH, et al. (2017) The challenge of mapping the human connectome based on diffusion tractography. *Nature Communications* 8(1):1349.

33. Yeh F-C, Verstynen TD, Wang Y, Fernández-Miranda JC, Tseng W-YI (2013) Deterministic Diffusion Fiber Tracking Improved by Quantitative Anisotropy. *PLOS ONE* 8(11):e80713.

34. Mišić B, et al. (2016) Network-Level Structure-Function Relationships in Human Neocortex. *Cereb Cortex* 26(7):3285–3296.

35. Crofts JJ, Higham DJ (2009) A weighted communicability measure applied to complex brain networks. *J R Soc Interface* 6(33):411–414.

36. Satterthwaite TD, et al. (2013) Functional Maturation of the Executive System during Adolescence. *J Neurosci* 33(41):16249–16261.

37. Ragland JD, et al. (2002) Working memory for complex figures: an fMRI comparison of letter and fractal n-back tasks. *Neuropsychology* 16(3):370–379.

38. Schlaggar BL, et al. (2002) Functional Neuroanatomical Differences Between Adults and School-Age Children in the Processing of Single Words. *Science* 296(5572):1476–1479.

39. Brown TT, et al. (2005) Developmental Changes in Human Cerebral Functional Organization for Word Generation. *Cereb Cortex* 15(3):275–290.

40. Ciric R, et al. (2017) Benchmarking of participant-level confound regression strategies for the control of motion artifact in studies of functional connectivity. *NeuroImage* 154:174–187.

41. Ciric R, et al. (2018) Mitigating head motion artifact in functional connectivity MRI. *Nat Protoc* 13(12):2801–2826.

42. Greve DN, Fischl B (2009) Accurate and robust brain image alignment using boundary-based registration. *Neuroimage* 48(1):63–72.

43. Hallquist MN, Hwang K, Luna B (2013) The nuisance of nuisance regression: spectral misspecification in a common approach to resting-state fMRI

preprocessing reintroduces noise and obscures functional connectivity. *Neuroimage* 82:208–225.

44. Thomas Yeo BT, et al. (2011) The organization of the human cerebral cortex estimated by intrinsic functional connectivity. *J Neurophysiol* 106(3):1125–1165.

45. Collin G, Scholtens LH, Kahn RS, Hillegers MHJ, van den Heuvel MP (2017) Affected Anatomical Rich Club and Structural–Functional Coupling in Young Offspring of Schizophrenia and Bipolar Disorder Patients. *Biological Psychiatry*. doi:10.1016/j.biopsych.2017.06.013.

46. Fukushima M, et al. (2017) Structure–function relationships during segregated and integrated network states of human brain functional connectivity. *Brain Struct Funct*:1–16.

47. Hill J, et al. (2010) Similar patterns of cortical expansion during human development and evolution. *PNAS* 107(29):13135–13140.

48. Margulies DS, et al. (2016) Situating the default-mode network along a principal gradient of macroscale cortical organization. *PNAS* 113(44):12574–12579.

49. Alexander-Bloch AF, et al. (2018) On testing for spatial correspondence between maps of human brain structure and function. *NeuroImage* 178:540–551.

50. Wood SN (2004) Stable and Efficient Multiple Smoothing Parameter Estimation for Generalized Additive Models. *Journal of the American Statistical Association* 99(467):673–686.

51. Wood SN (2011) Fast stable restricted maximum likelihood and marginal likelihood estimation of semiparametric generalized linear models. *Journal of the Royal Statistical Society: Series B (Statistical Methodology)* 73(1):3–36.

52. Preacher KJ, Hayes AF (2008) Asymptotic and resampling strategies for assessing and comparing indirect effects in multiple mediator models. *Behav Res Methods* 40(3):879–91.

53. Stiso J, Bassett DS (2018) Spatial Embedding Imposes Constraints on Neuronal Network Architectures. *Trends in Cognitive Sciences* 22(12):1127–1142.

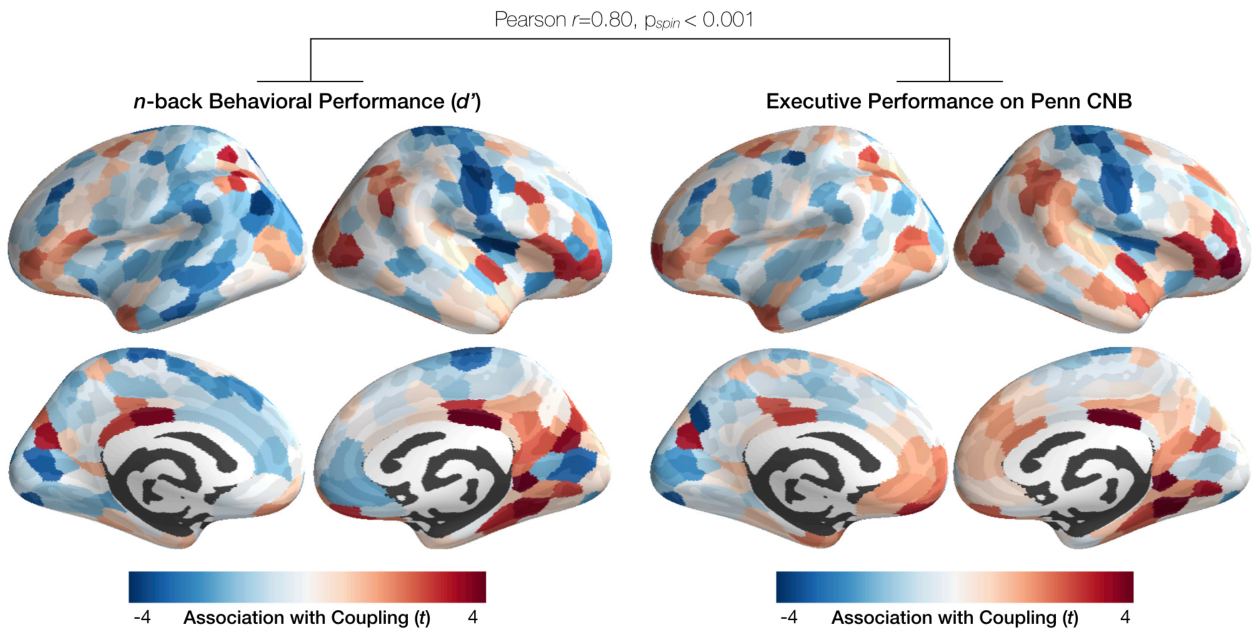


**Supplementary Figure 1.** *Regional coupling is similarly associated with n-back task performance and executive performance on a computerized battery*. We found that associations between regional structure-function coupling executive performance were consistent across two measures of performance. Variability in structure-function coupling was similarly associated with individual differences in performance on the n-back working memory task (*d'*), and a factor score summarizing accuracy on executive tasks administered as part of a separate computerized battery.

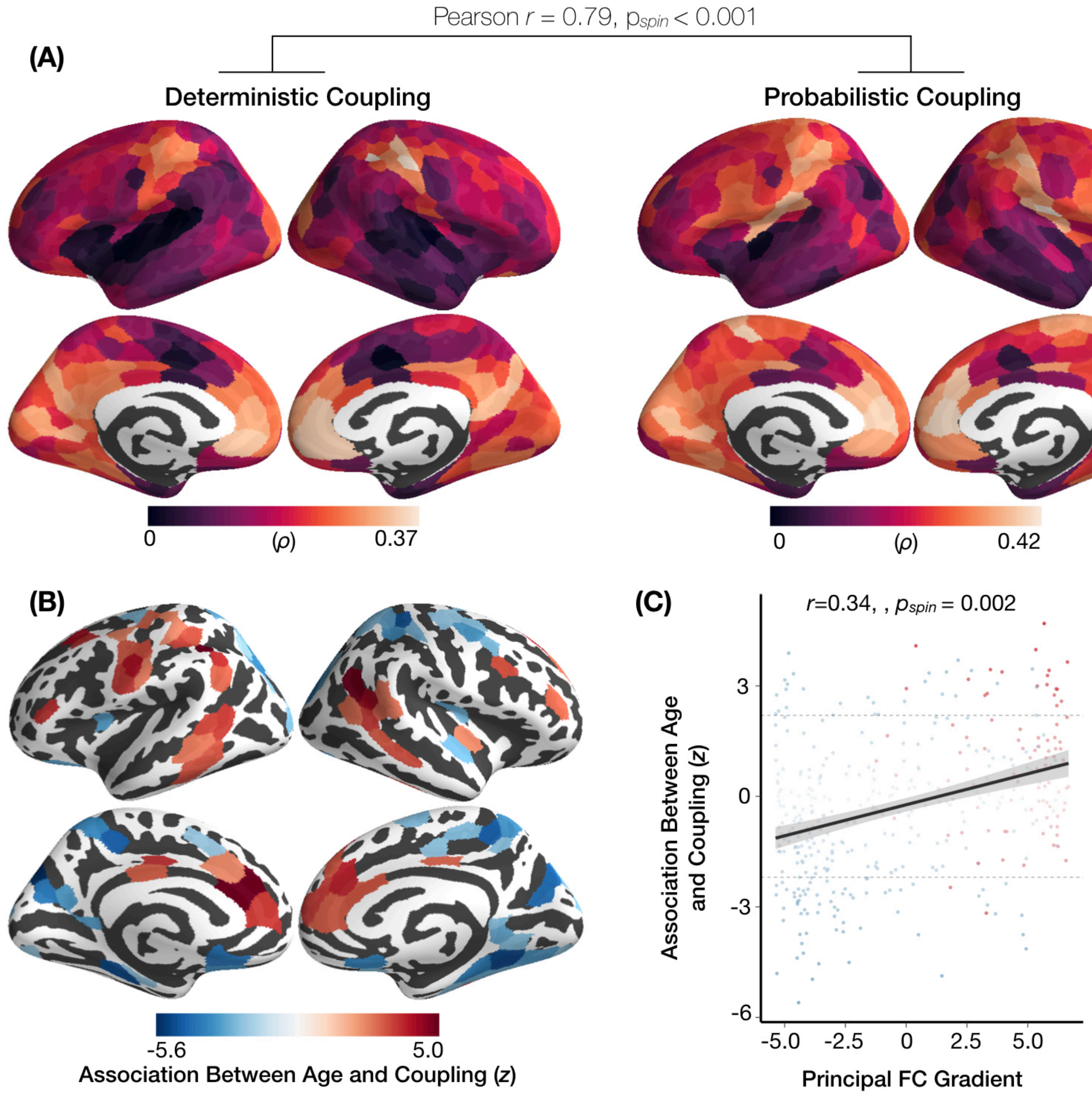


**Supplementary Figure 2.** *Hierarchy-dependent development of structure-function coupling using deterministic tractography.* Structural brain networks were derived from deterministic tractography and structural connection strength was modeled using communicability: a measure of inter-regional communication capacity that accounts for the strength of both direct and indirect structural connections between nodes. (**A**) Mean regional structure-function coupling was highly similar when calculated using deterministic or probabilistic tractography methods for constructing structural brain networks. (**B**) Age-related differences in structure-function coupling also remained highly consistent, and were distributed across superior temporal, parietal, cingulate, and prefrontal areas. (**C**) We observed hierarchy-dependent development of structure-function coupling: age-related increases in coupling were localized within transmodal areas of fronto-parietal and default networks, while age-related decreases in coupling were localized primarily within unimodal sensory areas. Multiple comparisons were controlled using the False Discovery Rate $Q<0.05$). Red points in panel C correspond to brain regions in the default mode network, while blue points represent regions in other functional systems. The significance of regional correlations was evaluated using non-parametric spatial permutation testing (denoted $p_{spin}$).

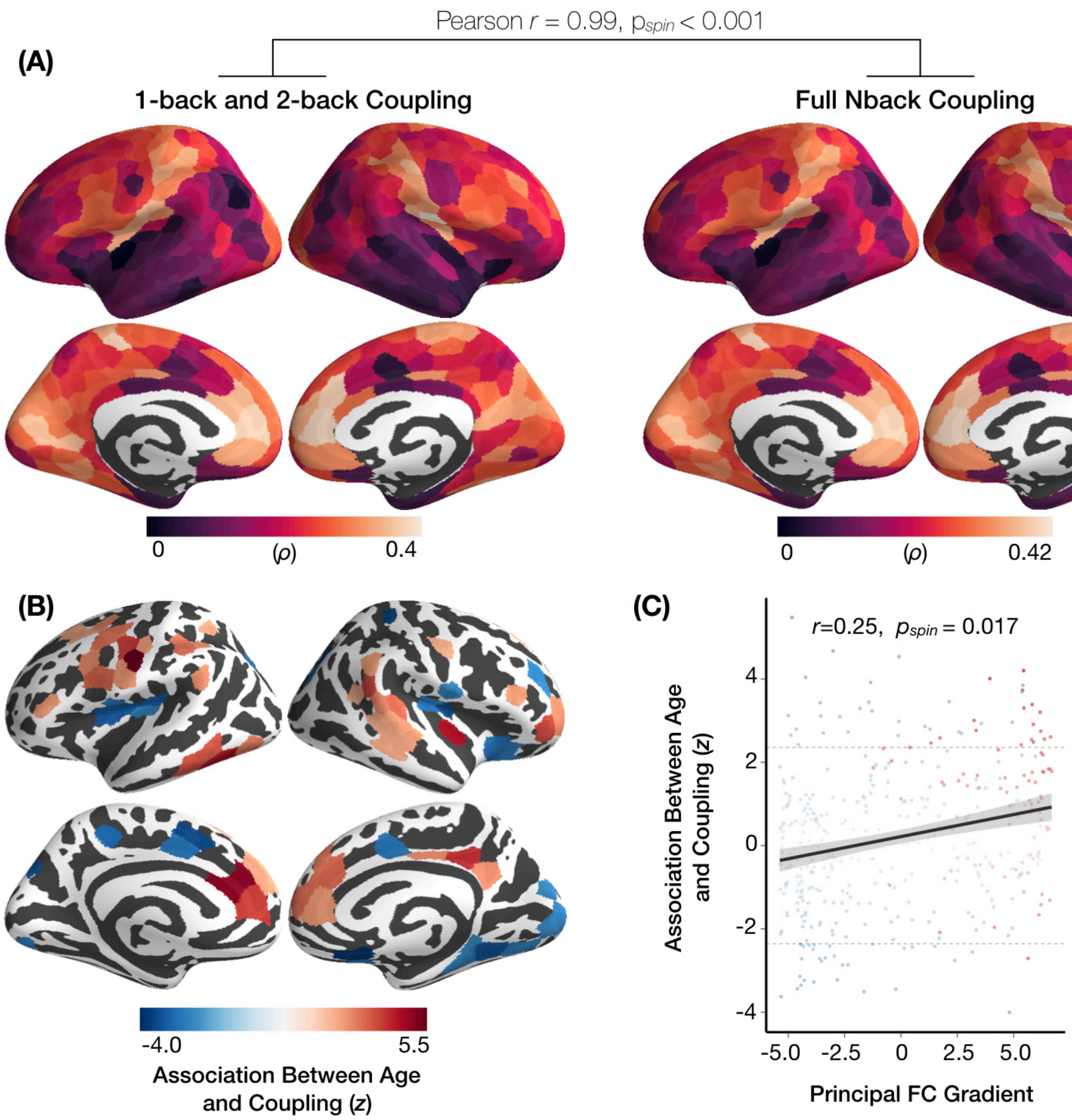


**Supplementary Figure 3.** *Hierarchy-dependent development of structure-function coupling estimating functional connectivity only during 1-back and 2-back task blocks.* Functional connectivity (FC) was estimated as the Pearson correlation coefficient between mean regional BOLD time-series during 1-back and 2-back blocks from the *n*-back working memory task. Structure-function coupling was then quantified using this working memory-related FC and structural connectivity derived from probabilistic tractography. (**A**) Mean regional structure-function coupling was highly similar when calculated using the full task time-series or high-load blocks of the *n*-back task. (**B**) Age-related changes in structure-function coupling also remained highly consistent, and were distributed across superior temporal, parietal, cingulate, and prefrontal cortex. (**C**) Consistent with other methodological approaches, we observed hierarchy-dependent development of structure-function coupling. Specifically, age-related increases in coupling were localized within transmodal areas of fronto-parietal and default networks, while age-related decreases in coupling were localized primarily within unimodal sensory areas. Multiple comparisons were controlled using the False Discovery Rate $Q<0.05$).Red points in panel C correspond to brain regions in the default mode network, while blue points represent regions in other functional systems. The significance of regional correlations was evaluated using non-parametric spatial permutation testing (denoted $p_{spin}$).

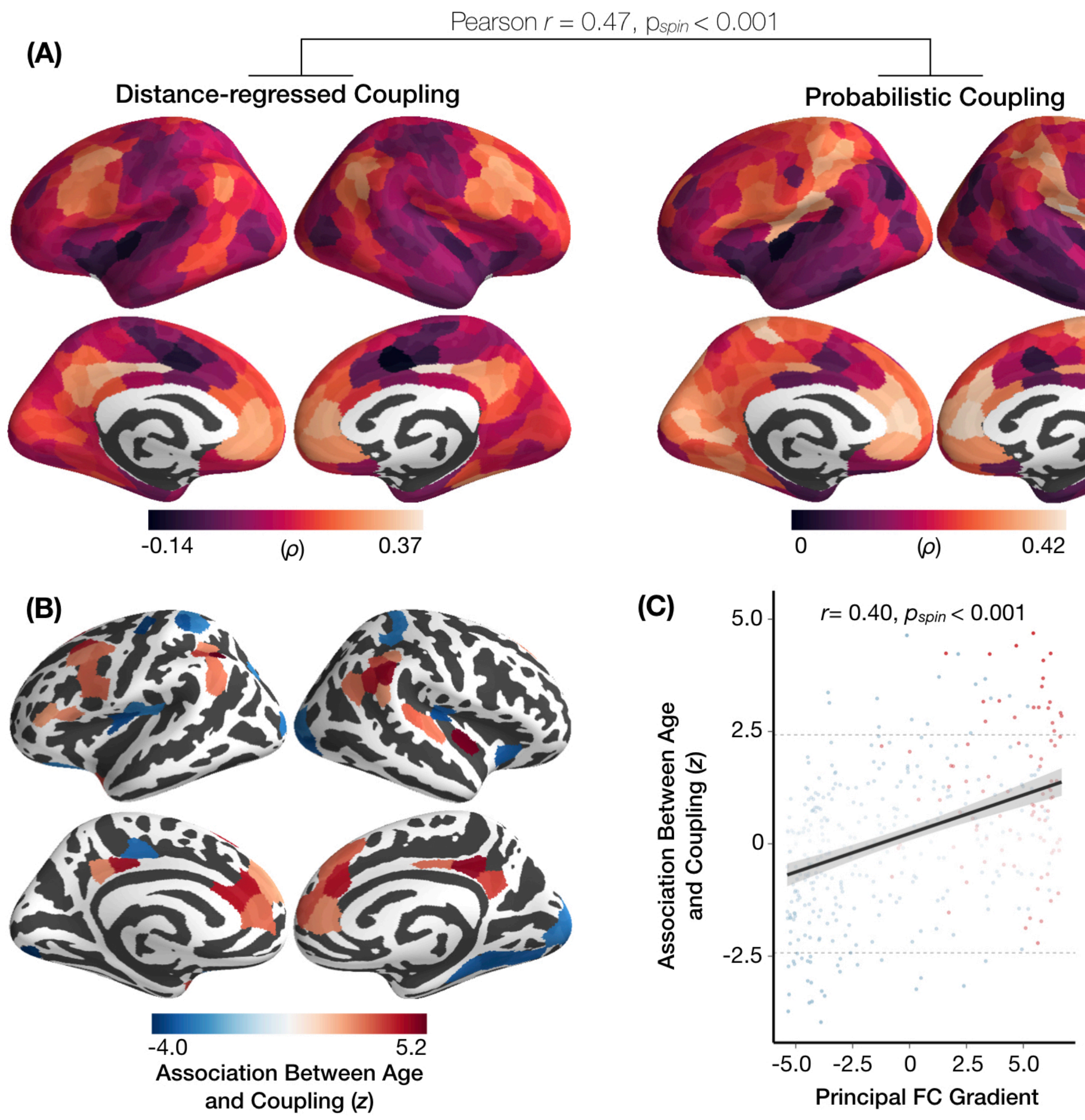


**Supplementary Figure 4.** *Hierarchy-dependent development of structure-function coupling when accounting for inter-regional distance.* Structure-function coupling was quantified as the partial correlation between regional structural and functional connectivity profiles while accounting for the Euclidean distance between brain regions. (**A**) Mean regional structure-function coupling aligned significantly with the coupling measures that did not account for inter-regional distance. Notably however, transmodal regions in frontoparietal and default networks exhibited higher structure-function coupling when accounting for the influence of inter-regional distance on coupling, while unimodal sensory regions had relatively lower structure-function coupling. (**B**) Age-related changes in structure-function coupling still remained highly consistent when accounting for inter-regional distance, and were distributed primarily in parietal, cingulate, and prefrontal cortex. (**C**) Consistent with other methodological approaches, we observed hierarchy-dependent development of structure-function coupling. Specifically, age-related increases in coupling were localized within transmodal areas of fronto-parietal and default networks, while age-related decreases in coupling were localized primarily within unimodal sensory areas. Multiple comparisons were controlled using the False Discovery Rate $Q<0.05$). Red points in panel C correspond to brain regions in the default mode network, while blue points represent regions in other functional systems. The significance of regional correlations was evaluated using non-parametric spatial permutation testing (denoted $p_{spin}$).

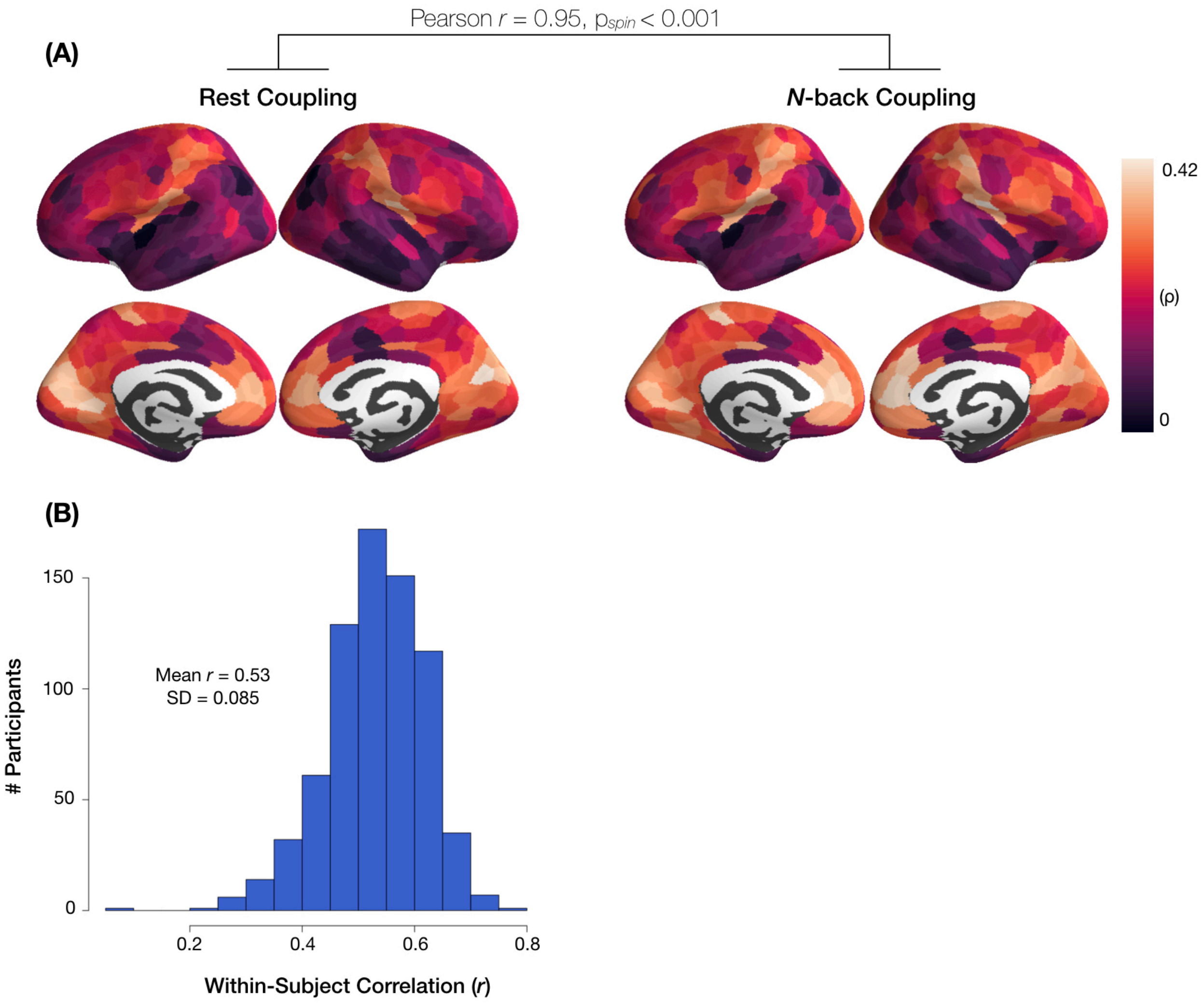


**Supplementary Figure 5.** *Global similarity and intra-individual changes in structure-function coupling between n-back and rest.* (**A**) When averaging across individuals, we found that spatial variability in mean structure-function coupling was highly consistent when using resting-state functional connectivity. (**B**) Within-subject correlations between structure-function coupling during *n*-back and rest reveal a greater degree of intra-individual variability in structure-function coupling during rest and task states.
